# Genome-wide identification of genes regulating DNA methylation using genetic anchors for causal inference

**DOI:** 10.1101/823807

**Authors:** Paul J. Hop, René Luijk, Lucia Daxinger, Maarten van Iterson, Koen F. Dekkers, Rick Jansen, BIOS Consortium, Joyce B.J. van Meurs, Peter A.C. ’t Hoen, M. Arfan Ikram, Marleen M.J. van Greevenbroek, Dorret I. Boomsma, P. Eline Slagboom, Jan H. Veldink, Erik W. van Zwet, Bastiaan T. Heijmans

**Affiliations:** Molecular Epidemiology, Department of Biomedical Data Sciences, Leiden University Medical Center, Leiden, 2333 ZC, The Netherlands; Department of Neurology, UMC Utrecht Brain Center, University Medical Centre Utrecht, Utrecht University, Utrecht 3584 CG, Netherlands; Department of Human Genetics, Leiden University Medical Center, Leiden, 2333 ZC, The Netherlands; Department of Psychiatry, Amsterdam UMC, Vrije Universiteit Amsterdam, Amsterdam Neuroscience, Amsterdam, 1081 HV, The Netherlands; Biobank-based Integrated Omics Study Consortium. For a complete list of authors, see the acknowledgements.; Department of Internal Medicine, Erasmus Medical Centre, Rotterdam, The Netherlands; Centre for Molecular and Biomolecular Informatics, Radboud Institute for Molecular Life Sciences, Radboud University Medical Center Nijmegen, Nijmegen, The Netherlands; Department of Epidemiology, Erasmus University Medical Center, Rotterdam, 3015 CE, The Netherlands; Department of Internal Medicine, Maastricht University Medical Center, Maastricht, 6211 LK, The Netherlands; School for Cardiovascular Diseases (CARIM), Maastricht University Medical Center, Maastricht, 6229 ER, The Netherlands; Department of Biological Psychology, Vrije Universiteit Amsterdam, Neuroscience Campus Amsterdam, Amsterdam, 1081 BT, The Netherlands; Medical Statistics, Department of Biomedical Data Sciences, Leiden University Medical Center, Leiden, Zuid-Holland 2333 ZC, The Netherlands

## Abstract

DNA methylation is a key epigenetic modification in human development and disease, yet there is limited understanding of its highly coordinated regulation. Here, we identified 818 genes that influence DNA methylation patterns in blood using large-scale population genomics data. By employing genetic instruments as causal anchors, we identified directed associations between gene expression and distant DNA methylation levels, whilst ensuring specificity of the associations by correcting for linkage disequilibrium and pleiotropy among neighboring genes. We found that DNA methylation patterns are commonly shaped by transcription factors that consistently increase or decrease DNA methylation levels. However, we also observed genes encoding proteins without DNA binding activity with widespread effects on DNA methylation (e.g. *NFKBIE, CDCA7(L)* and *NLRC5*) and we suggest plausible mechanisms underlying these findings. Many of the reported genes were unknown to influence DNA methylation, resulting in a comprehensive resource providing insights in the principles underlying epigenetic regulation.

## INTRODUCTION

The epigenome is fundamental to development and cell differentiation. Dysregulation of the epigenome is a hallmark of many diseases, ranging from rare developmental disorders to common complex diseases and aging (Bjornsson, 2015; Dor and Cedar, 2018; Sen et al., 2016). The epigenome is highly dynamic and is extensively modified and remodeled in response to external and internal stimuli (Zentner and Henikoff, 2013). However, the networks underlying these highly coordinated epigenetic modifications remain to be fully elucidated. Hence, the systematic identification of genes that are involved in epigenetic regulation and the determination of their respective target sites will be a key element towards an in-depth understanding of epigenomic (dys)regulation.

DNA methylation is a key component of the epigenome that controls, stabilizes and/or marks the transcriptional potential of a genomic region (Schübeler, 2015). It involves the addition of a methyl group onto cytosines, mainly at CpG dinucleotides. Although considerable research has been devoted to studying the enzymes that *write, maintain* and *erase* DNA methylation (i.e., DNMTs and TETs) (Medvedeva et al., 2015), less is known about factors that are otherwise involved in the regulation of DNA methylation. These may include proteins and non-coding RNAs that regulate, interact with or recruit the DNA methylation machinery (Shen and Laird, 2013). Transcription factors, for example, do not only act indirectly by regulating the transcription of epigenetic genes, but have also been shown to control the DNA methylation state of their target sites by recruiting or repelling DNMT or TET proteins (Blattler and Farnham, 2013; Marchal and Miotto, 2015). Experimental evidence for genes involved in the regulation of DNA methylation has been mainly obtained from *in vitro* experiments focusing on single genes or is based on animal models (Daxinger et al., 2013; Marchal and Miotto, 2015; Stadler et al., 2011; Wang et al., 2015). A comprehensive genome-wide resource of genes affecting DNA methylation in humans is currently lacking.

We recently developed a method to identify directed and specific gene-gene interactions in population omics data (Luijk et al., 2018). Instead of using measured gene expression, this method builds upon previous work in which genetic variants were utilized as causal anchors for gene expression (Gamazon et al., 2015; Gusev et al., 2016). This allows for the identification of directed and unconfounded associations within observational data. Here, we adapt this method to establish a resource of 818 genes that influence DNA methylation using genomic, methylomic and transcriptomic data in up to 4,056 individuals (Bonder et al., 2016; Zhernakova et al., 2016). We show that this resource reveals insight into the mechanisms underlying epigenetic regulation.

## RESULTS

### Establishing a resource of genes that influence DNA methylation

In order to identify genes that influence DNA methylation, we employed an approach that consists of two parts. First, we identified predictive genetic variants for the expression of each gene in our data, which we aggregated into single predictive scores termed genetic instruments (GIs) (Luijk et al., 2018). Second, we used these GIs as causal anchors to establish *directed* effects of gene expression on genome-wide DNA methylation levels while ensuring that these associations were *specific* by accounting for linkage disequilibrium (LD) and pleiotropy among neighboring GIs (see figure 1 for an overview of the successive steps in the analysis).

**Figure 1.**
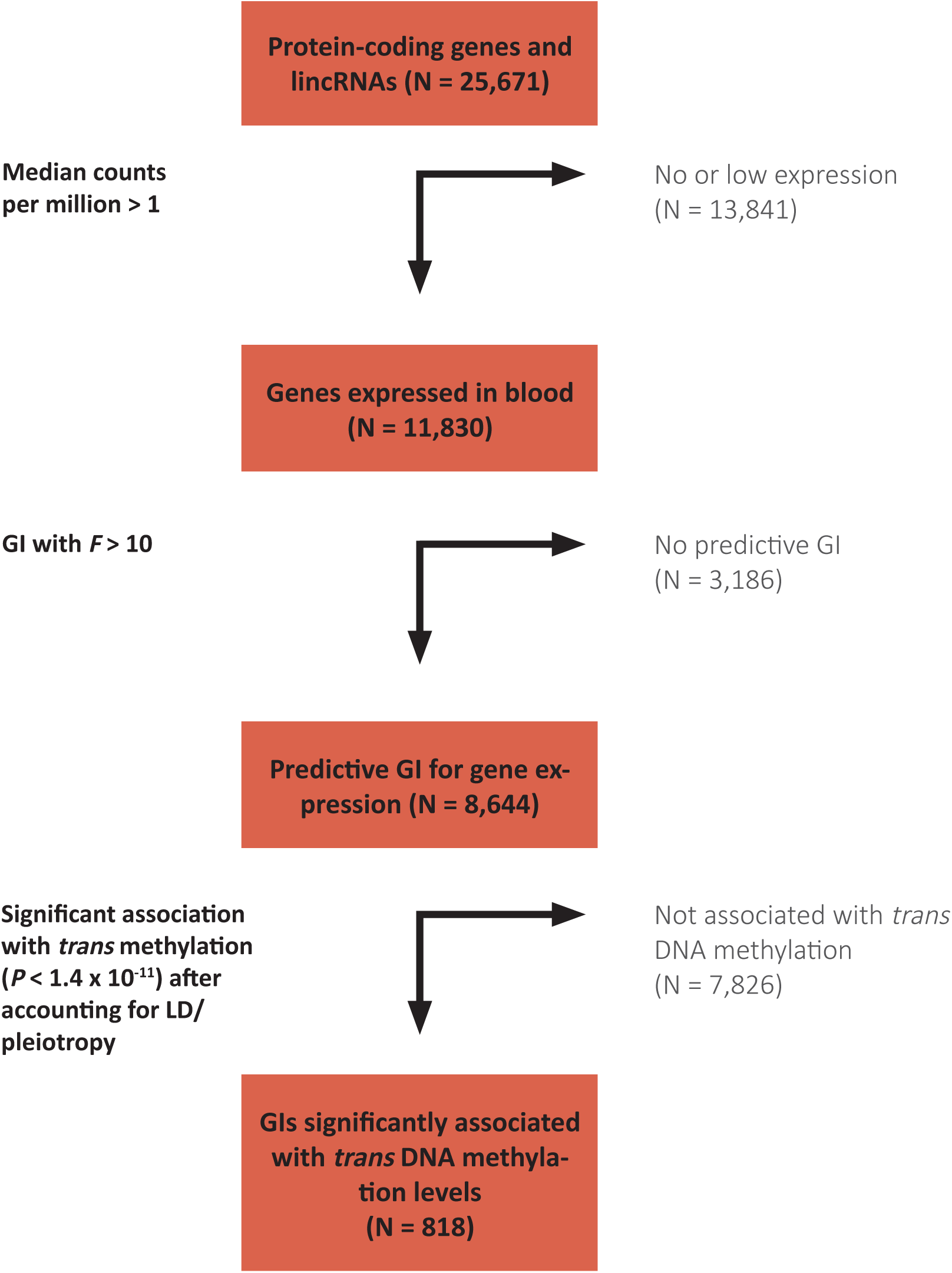
Flowchart showing the successive steps leading to the resource of 818 genes that influence DNA methylation *in trans*.

To construct the genetic instruments, we used data on 3,357 unrelated individuals with available genotype and RNAseq data derived from whole-blood. We focused the analysis on 11,830 expressed genes (median counts per million > 1). In the training set (1/3 of the data, 1,119 individuals) we applied penalized regression (LASSO) to the expression of each gene to obtain a GI that consists of 1 or more SNPs aggregated into a single score that is predictive of expression (Tibshirani, 1996). Here, we corrected the expression data for age, sex, biobank, blood cell composition and five principal components. We then assessed the predictive ability of the constructed GIs in a separate test set of 2,238 individuals by predicting their gene expression values using these GIs. Of the 11,830 tested GIs, 8,644 were sufficiently predictive of expression levels in the test set to serve as valid genetic instruments (*F*-statistic > 10, median R^2^ = 0.04, Table S1) (Staiger and Stock, 1997).

Next, we tested for an association between all 8,644 predictive GIs and genome-wide DNA methylation levels at 428,126 autosomal CpG-sites *in trans* (>10Mb distance from the tested gene), using genotype and DNA methylation data (Illumina 450k array) derived from whole blood of 4,056 unrelated individuals (3,251 samples overlap with RNAseq data). These associations were computed using linear regression, while correcting for age, sex, blood cell composition, biobank, five principal components, and corrected for bias and inflation in the test-statistics (van Iterson et al., 2017). These analyses resulted in *directed* associations between 2,223 genes and 5,284 CpGs (Bonferroni correction, *P* < 1.4 × 10^−11^; Table S2). Although directed, the associations resulting from this analysis may not be specific for a single gene as linkage disequilibrium (LD) and/or pleiotropy may result in GIs that are predictive of multiple neighboring genes (Luijk et al., 2018). We therefore adjusted all significant GI-CpG pairs for all neighboring GIs (<1Mb) to account for correlation induced by LD/pleiotropy among neighboring genes in order to identify the specific gene in a region driving the association. Next, we removed genes with potential residual pleiotropic effects on the expression of neighboring significant genes (*F* > 5) (together these two steps led to the removal of 1,387 genes and 2,844 CpGs; Table S3). Finally, we excluded effects of long-range pleiotropy and LD (by rerunning the analysis for CpGs influenced by multiple genes from the same chromosome, including all these genes in the model; removing 6 genes, 13 CpGs), and residual effects of white blood cell composition (by correcting for genetic variants known to be associated with WBC; removing 12 genes, 43 CpGs, figure S1) (Orrù et al., 2013; Roederer et al., 2015). The validity of these results was corroborated by a comparison with previous *trans*-methylation QTL studies in blood. Although not designed to infer genes that are specifically responsible for associations, such studies are expected to produce partly overlapping outcomes. We found that 692 target CpGs identified in our study were reported in two previous independent *trans*-meQTL studies (*P* < 1 × 10^−40^) (Gaunt et al., 2016; Lemire et al., 2015). For the great majority of overlapping CpGs, the corresponding GI and *trans*-meQTLs SNP were in close proximity (Table S4-6).

The final result of our step-wise analysis was a collection of 818 genes with directed and specific associations with DNA methylation levels of 2,384 unique target CpGs *in trans* (Bonferroni correction, *P* < 1.4 × 10^−11^; (Table S7).

### Function of genes that influence DNA methylation *in trans*

As shown in figure 2, a considerable fraction (N = 308) of the identified genes influenced multiple CpGs *in trans* (Table S8). We observed that for these genes, the direction of effect was often skewed towards either increased or decreased methylation levels at the target CpGs (Figure 2a). For 30 out of 37 genes that were associated with at least 10 CpGs, the direction of effect was significantly skewed towards increased (19 genes) or decreased (11 genes) methylation levels respectively (binomial test, FDR < 0.05, Table S9). We first considered two previously hypothesized molecular roles of the identified genes: transcription factors (Lambert et al., 2018) and core epigenetic factors (Medvedeva et al., 2015), which we will now consider in more detail.

**Figure 2.**
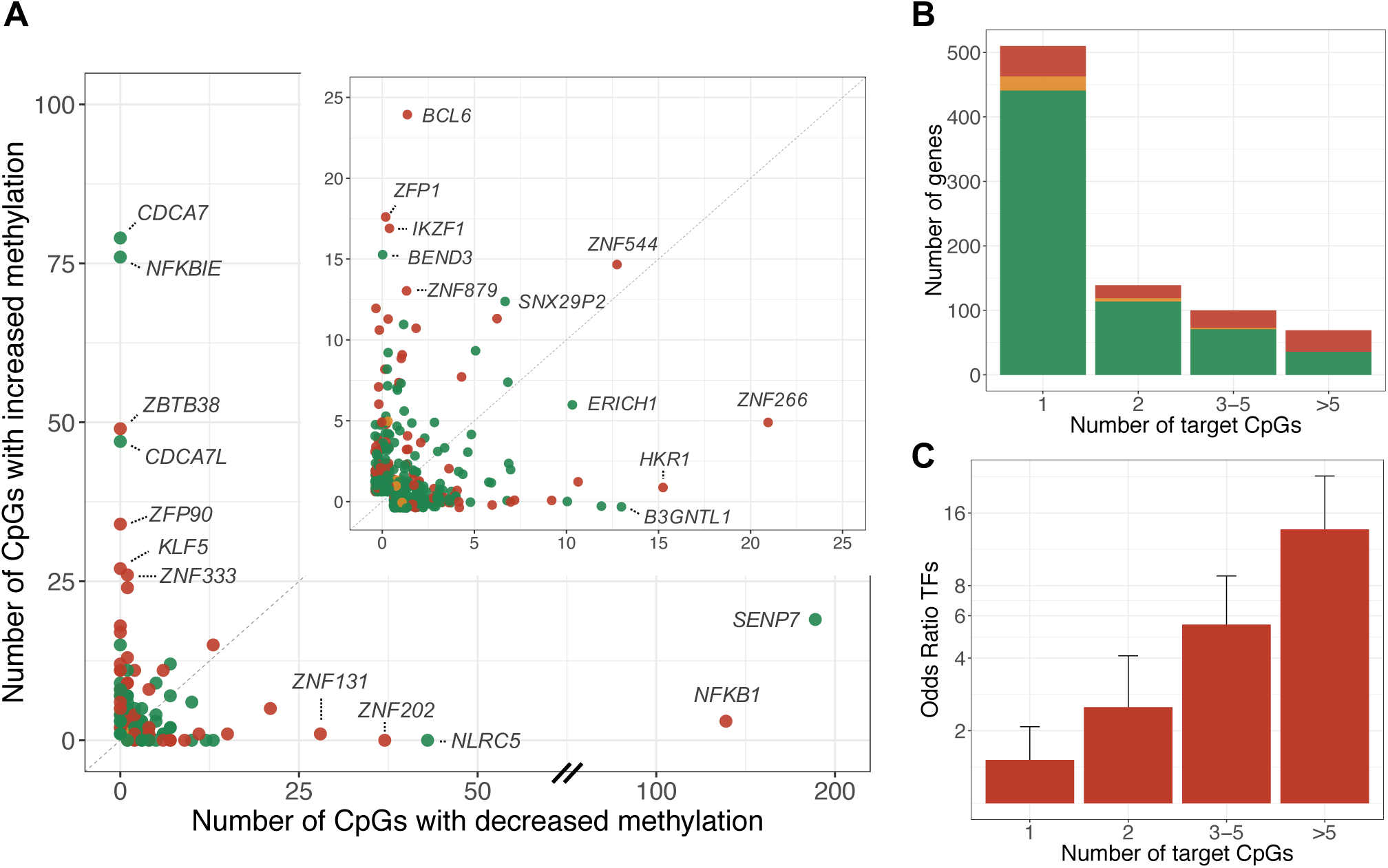
A considerable fraction of the identified genes (N = 308) influenced multiple target CpGs *in trans*. **(a)** Each dot represents a gene with *trans* DNA methylation effects. The x-axis shows the number of affected target CpGs with decreased methylation levels upon increased gene expression, the y-axis shows the number of affected target CpGs with increased methylation levels upon increased gene expression. The figure in the right upper corner is a zoomed-in version in which only genes that affect less than 25 CpG-sites in either direction are displayed. **(b)** Bars represent the number of genes with either 1, 2, 3-5 or more than 5 target CpGs. The percentage of genes that are annotated as transcription factors increases with the number of target CpGs. **(c)** Enrichment (odds ratio) for transcription factors for genes with either 1,2, 3-5 or more than 5 target CpGs. Error bars represent 95% confidence intervals.

#### Transcription factors

We found that the resource (818 genes) was enriched for transcription factors (N = 127, odds ratio = 2.74, *P* = 3.1 × 10^−18^) using a manually curated list of transcription factors (TFs) (Lambert et al., 2018). As shown in figure 2c this enrichment was driven by TFs that were associated with multiple target CpGs and there was a stronger TF enrichment with an increasing number of target CpGs. In total, 80 (63%) of the significant TFs in our data influenced more than 1 CpG-site. The direction of effect of the TFs on their target CpGs was either skewed towards increased or decreased methylation levels respectively (of the 23 TFs associated with at least 10 CpGs, 6 were significantly associated with lower methylation levels and 14 with higher methylation levels). Transcription factors influencing the most CpGs included *NFKB1*, a key immune regulator (142 target CpGs), *ZBTB38*, a methyl-binding TF (49 target CpGs) and *ZNF202*, a zinc finger protein involved in lipid metabolism (37 target CpGs) (Filion et al., 2006; Hayden et al., 2006; Wagner et al., 2000). We found that 100 out of the 127 (79%) transcription factors belonged to the C2H2 Zinc Finger family (odds ratio = 3.07, *P* = 5.2 × 10^−7^), of which the majority (N = 70) contains a KRAB-domain. In line with the enrichment for TFs and Zinc Fingers, we found that the gene set was overrepresented in the GO terms Nucleic Acid binding (N = 99, *P* = 1.1 × 10^−14^), DNA Binding (N = 114, *P* = 4.7 × 10^−9^), Metal Ion binding (N = 146, *P* = 1.4 × 10^−8^) and transcription factor activity (N = 73, *P =* 4.4 × 10^−8^) (Table S10).

#### Core epigenetic factors

Next, we compared our findings with a manually curated database of core epigenetic factors (EpiFactors) (Medvedeva et al., 2015). This database is mainly focused on the core enzymes that directly write/maintain/establish epigenetic marks, but it does also include a few ‘borderline cases’ such as TFs that interact with epigenetic proteins. We found that 36 of the identified genes overlapped with genes in this database, which did not constitute an enrichment (odds ratio = 1.02, *P* > 0.05). Interestingly, however, the majority of the 36 genes encode proteins that target histone proteins (26 out of 36). In addition, 7 genes were annotated as transcription factors in the manually curated TF catalog (Lambert et al., 2018). The core epigenetic factor associated with most target CpGs include transcription factor *IKZF1* (positively associated with methylation at 17 target CpGs), histone demethylase *KDM5B* (positively associated with methylation at 7 target CpGs), and *BRD3*, which recognizes acetylated lysine residues on histones (positively associated with methylation at 5 target CpGs). The significant core epigenetic factors also included the DNA methyltransferase *DNMT3A*, which was associated with increased methylation at five target CpGs. Further exploration of potential *DNMT3A* targets indicated that the test-statistics of *DNMT3A* were skewed towards increased DNA methylation levels, compatible with widespread but small effects (figure S2A). Of note, of the other main DNA methylation modifiers (*DNMT1, DNMT3B, TET1,2,3*) we had a sufficiently predictive GI for *DNMT1* only (*F =* 90, *R*^*2*^ = 0.04). However, we did not find significant associations for this gene (figure S2B).

#### Other mechanisms underlying regulation of DNA methylation

Finally, the majority of the identified genes (N = 662), did not belong to the two *a priori* categories TFs and core epigenetic factors despite including genes that were associated with multiple CpG-sites (figure 2). A small fraction of these genes encodes proteins with DNA-binding properties (N = 24). *BEND3* for example, is a DNA-binding protein that was associated with increased methylation at 15 CpG-sites. Previous research has shown that BEND3 represses transcription by attracting the MBD3/NuRD complex that initiates histone deacetylation (Saksouk et al., 2014).

A plausible explanation for the identification of non-DNA binding proteins is that they affect DNA methylation indirectly through interaction with epigenetic proteins. This is illustrated by several examples of genes that encode non-DNA-binding proteins.

#### NFKBIE

The *NFKBIE* gene encodes IκBε which is an inhibitor of NFκB, a transcription factor that plays a fundamental role in the regulation of the immune response (Bonizzi and Karin, 2004; Liu et al., 2017). IκBε binds to components of NFκB and retains it in the cytoplasm, thereby preventing it from activating genes in the nucleus. Consistent with the previous interpretation of a *trans*-methylation QTL effect (Bonder et al., 2016), we found that increased expression of *NFKB1* was associated with genome-wide loss of DNA methylation. In contrast, we found that increased expression of *NFKBIE* resulted in *higher* methylation levels at 76 CpG-sites across the genome. In line with its role as NFκB inhibitor, a substantial number of its target CpGs (28) overlap with NFκB’s target CpGs and show opposite effects (figure 3a).

**Figure 3.**
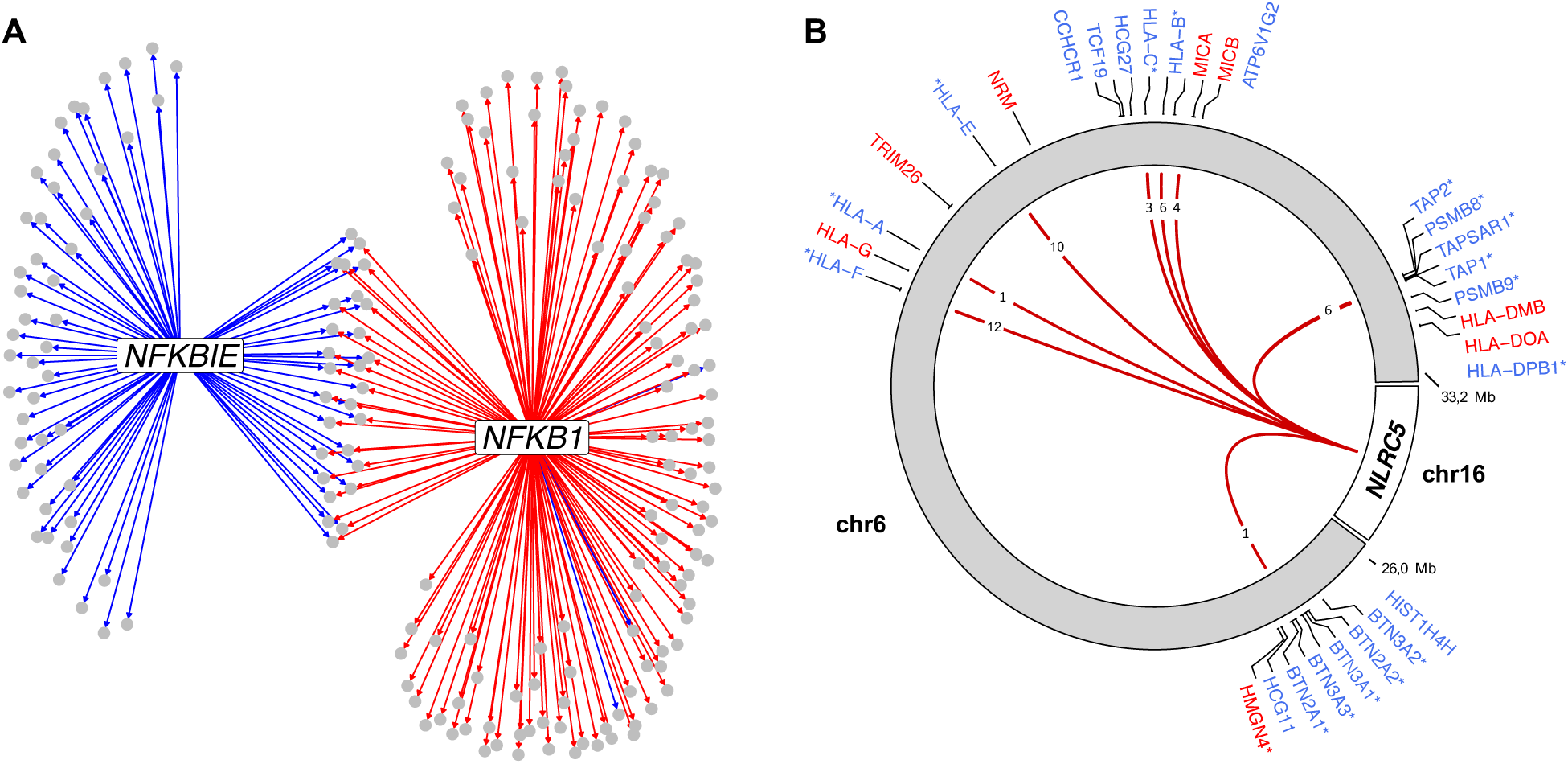
**(a)** Network for transcription factor *NFKB1* and its inhibitor *NFKBIE*. Grey circles indicate target CpGs and arrows represent directed associations (i.e. association between GI and DNA methylation levels). Blue lines indicate a positive association between gene expression and DNA methylation levels, red lines indicate a negative association between gene expression levels and DNA methylation levels. **(b)** *NLRC5* (chromosome 16) was associated with decreased DNA methylation levels at multiple (N = 43) CpG-sites in the classical and extended MHC-region (chromosome 6). Red lines indicate a negative association between *NLRC5* expression levels and DNA methylation levels. The numbers displayed in the lines indicate how many target CpGs the line represents. Gene labels are displayed if one or more target CpGs were associated with the expression of these genes. Blue gene symbols refer to genes negatively correlated with target CpG methylation (implying upregulation by *NLRC5*), and vice versa for red labels. Asterisks indicate that the GI corresponding to *NLRC5* was also (positively) associated with this gene.

#### NLRC5

Increased expression of *NLRC5* was associated with decreased methylation levels at 43 CpG-sites that were all located in either the classical or the extended MHC-region (Mungall et al., 2003). NLRC5 is a known activator of MHC class I genes, and in line with this we found that the methylation levels of most target CpGs (N = 36) were negatively associated with the expression levels of one or more neighboring MHC-genes (Figure 3b/Table S11-12). Furthermore, the GI corresponding to *NLRC5* was positively associated with 16 of these genes. NLRC5 itself does not contain a DNA-binding domain, instead it has been shown to affect transcription by cooperating with a multi-protein complex that is assembled on the MHC class I promoter (Kobayashi and van den Elsen, 2012). Interestingly, NLRC5 acts as a platform for enzymes that open chromatin by histone acetylation and/or demethylation of histone H3, indicating that decreased DNA methylation may be a consequence of altered chromatin state.

#### SENP7

The gene with the largest number of detected target CpGs was *SENP7*. It was associated with decreased methylation levels at 189 target CpGs, and with increased methylation levels at 19 target CpGs. The majority (86%) of the target CpGs were located on the q-arm of chromosome 19. We found that for most of these CpGs (92%) their DNA methylation levels were associated with the expression levels of one or more nearby zinc fingers (table S13-14), consistent with previous gene network analyses (Luijk et al., 2018). Although SENP7 has no DNA-binding properties, previous research has shown that it exerts its effect through deSUMOylation of the chromatin repressor KAP1 (Garvin et al., 2013). KAP1 can act as a scaffold for various heterochromatin-inducing factors and there is emerging evidence that KAP1 is directly involved in regulating DNA methylation (Quenneville et al., 2011; Zuo et al., 2012). SENP7 is therefore likely to affect DNA methylation through its interaction with KAP1.

#### CDCA7

Mutations in *CDCA7* have been shown to cause ICF, a rare primary immunodeficiency characterized by epigenetic abnormalities (Thijssen et al., 2015). Previous research showed that *CDCA7*-mutated ICF patients show decreased DNA methylation levels at pericentromeric repeats and heterochromatin regions and, similarly, *CDCA7* depletion in mouse embryonic fibroblasts leads to decreased DNA methylation at centromeric repeats (Thijssen et al., 2015; Velasco et al., 2018). In line with this we found that increased expression of *CDCA7* is associated with increased methylation levels at 79 CpG-sites that were distributed across chromosomes (figure 4a) and were enriched in low-activity regions (e.g. quiescent states; figure 4b) and repeat sequences (odds ratio 2.13, *P* = 0.006). In addition, a volcano plot showed that the test-statistics of *CDCA7* were highly skewed towards positive effects, suggesting that *CDCA7* has widespread effects on DNA methylation (figure S3a).

**Figure 4.**
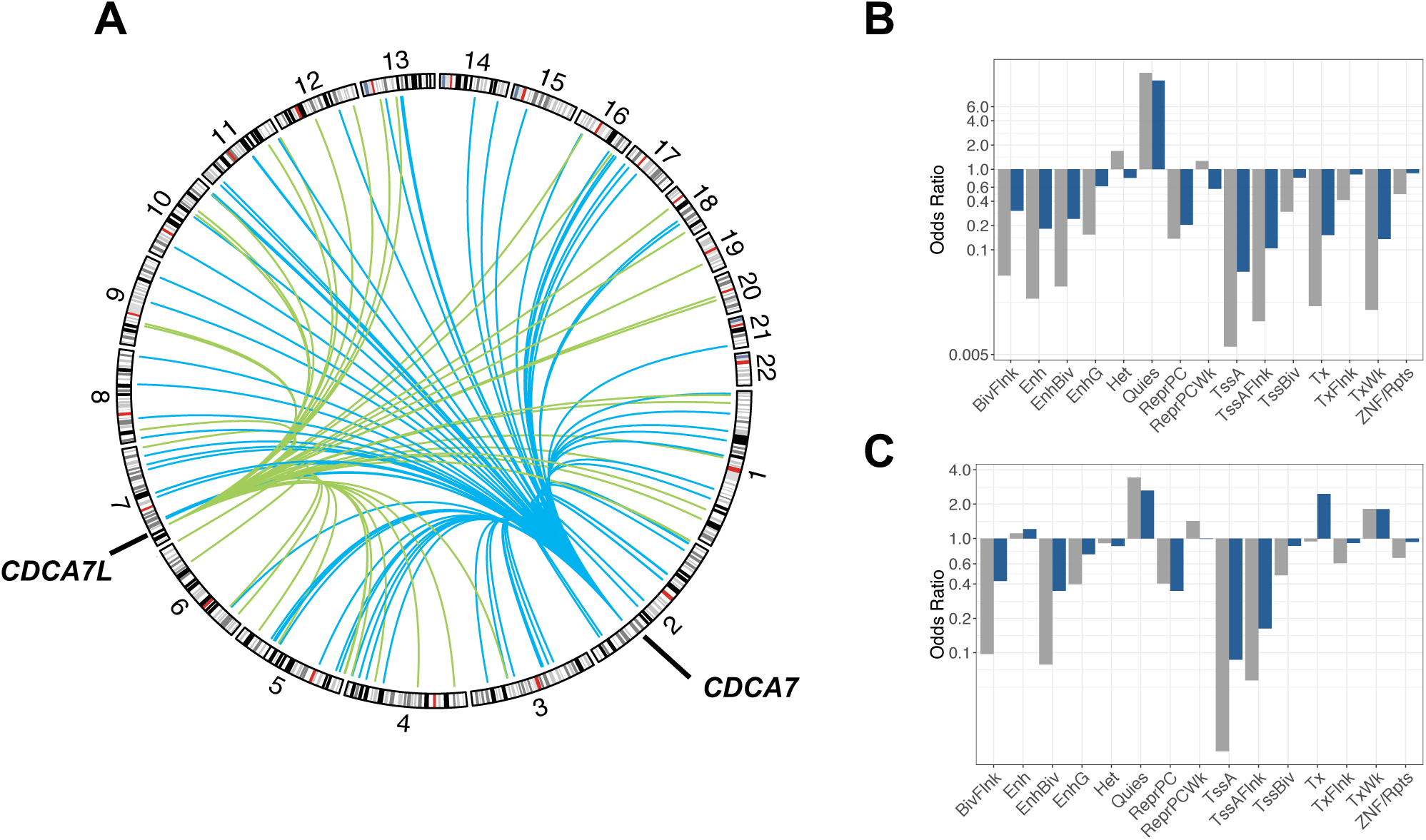
**(a)** *CDCA7* (located on chromosome 2) and *CDCA7L* (located on chromosome 7) both influence genome-wide DNA methylation levels. Blue lines indicate a positive association between *CDCA7* expression and *trans* DNA methylation levels. Green lines indicate a positive association between *CDCA7L* expression levels and *trans* DNA methylation levels. **(b**,**c)** Over- or underrepresentation of target CpGs in predicted chromatin states for **(b)** *CDCA7* and **(c)** *CDCA7L*. Blue bars represent enrichment of CpGs that are significant at a genome-wide significance level (*P* < 1.4 × 10^−11^) and grey bars represent enrichment of CpGs that are significant at a gene-level significance level (*P* < 1.2 × 10^7^). BivFlnk: flanking bivalent TSS/enhancer, Enh: enhancer, EnhBiv: bivalent enhancer, EnhG: genic enhancer, Het: heterochromatin, Quies: quiescent, ReprPC: repressed polycomb, ReprPCWk: weak repressed polycomb, TssA: active TSS, TssAFlnk: flanking active TSS, TssBiv: bivalent/poised TSS, Tx: strong transcription, TxFlnk: weak transcription, ZNF/Rpts: ZNF genes & repeats.

#### CDCA7L

*CDCA7L* is a paralog of *CDCA7* and, similarly, its increased expression was associated with a genome-wide increase of DNA methylation levels (47 CpG-sites; figure 4a/figure S3b). *CDCA7L*’s target CpGs did not overlap with those of *CDCA7*, however, they did show a similar genomic distribution and were enriched in inactive regions (Figure 4c), although enrichment at repeat regions was reduced (OR = 1.59, *P* > 0.05). Interestingly, previous research has shown that the risk allele of the genetic variant most highly associated with multiple myeloma (rs4487645) was associated with increased *CDCA7L* expression (Li et al., 2016). Our GI for *CDCA7L* consisted of 5 SNPs, of which one (rs17361667) was in strong LD (*r*^*2*^ = 0.7) with the risk variant rs4487645. If the risk variant indeed exerts its pathogenic effect through an effect on *CDCA7L* expression, *CDCA7L’s* effects on DNA methylation might be involved in the pathogenesis of multiple myeloma. Moreover, we found that our multi-SNP GI was a stronger predictor of *CDCA7L* expression (*F* = 171) as compared with rs4487645 (*F* = 60), and may therefore be useful in gaining more insight into the role of *CDCA7L* in multiple myeloma.

## DISCUSSION

Our genome-wide analysis, utilizing genetic instruments for gene expression, identified 818 genes that influence distant DNA methylation levels in blood and as such provides a novel resource that reveals insights into the principles of epigenetic regulation.

We found that DNA methylation levels are commonly influenced by transcription factors. The direction of effect is often consistent for each TF, and can either be skewed towards increased or decreased methylation levels at the target CpGs. In line with these findings, previous studies suggest that TFs can regulate the acquisition and/or loss of DNA methylation at their binding sites (Blattler and Farnham, 2013; Marchal and Miotto, 2015; Zhu et al., 2016). For example, several TFs have been shown to recruit DNMTs to their binding sites, thereby causing *de novo* DNA methylation (Brenner et al., 2005; Di Croce et al., 2002; de la Rica et al., 2013; Velasco et al., 2010). Conversely, TFs have been indicated to protect against the acquisition of DNA methylation by blocking *de novo* methylation or by interacting with TET proteins (Bonder et al., 2016; de la Rica et al., 2013; Stadler et al., 2011; Wang et al., 2015). We identified TFs with a previously unrecognized role in the regulation of DNA methylation (e.g. *ZNF202, ZNF131* and *ZFP90*), and provided support for the presumed role of TFs as previously implicated by post-hoc interpretation of results from methylation QTL mapping (*NFKB1* and *ZBTB38*) (Bonder et al., 2016). Interestingly, the identified TFs are overrepresented in the C2H2-zinc finger family, which is in line with previous *trans*-meQTL findings (Lemire et al., 2015) The majority of the identified TFs contain a Krüppel-associated box (KRAB) domain which has been implicated in epigenetic silencing through the recruitment of KAP1 to their binding sites. KAP1 subsequently recruits proteins that establish heterochromatin such as the NuRD-complex and possibly DNMTs, thereby causing *de novo* methylation (Groner et al., 2010; Iyengar et al., 2011; Meylan et al., 2011). Although we found 8 KRAB-ZFs with at least 10 target CpGs that were significantly skewed towards increased methylation, 4 were associated with decreased methylation. A possible explanation is that not all KRAB-ZFs act via KAP1. For example, the KRAB-ZF *ZNF202*, which was negatively associated with 37 target CpGs, contains a SCAN domain that prevents the binding of KAP1 (Lupo et al., 2013). Overall, our systematic genome-wide analysis identifies novel epigenetic regulatory functions for TFs, significantly expands upon TFs that were previously implicated in DNA methylation regulation, and identifies the direction of the effect on DNA methylation.

Exploration of the genes that do not encode transcription factors revealed several potential mechanisms through which these genes may influence DNA methylation. First, several of these genes encode proteins with DNA-binding properties, which might recruit or block the DNA methylation machinery in a similar way to TFs. *BEND3* for example, encodes a DNA-binding protein that attracts the chromatin remodeling NuRD complex to its binding sites (Saksouk et al., 2014). Second, closer inspection of proteins that do not have DNA-binding properties suggests that they may regulate DNA methylation through protein-protein interactions. Possible mechanisms include post-translational regulation (*NFKBIE* encodes for IκBε which retains NF-kB in the cytoplasm (Bonizzi and Karin, 2004)), post-translational modification (SENP7: deSUMOylates the repressor KAP1 (Luijk et al., 2018)) and recruitment of epigenetic proteins to specific target sites through association with a DNA-binding protein (NLRC5 associates with a protein complex in MHC-I region (Kobayashi and van den Elsen, 2012)). Third, a fraction of the identified genes overlap with genes in a database that focuses on the core epigenetic regulators (i.e. the main enzymes that write/erase epigenetic marks such as DNMTs and histone acetyltransferases) (Medvedeva et al., 2015). Interestingly, we had a predictive GI for the expression levels of *DNMT1* and *DNMT3a*, but found only 5 significant associations for *DNMT3A* and no significant associations for *DNMT1*. Although this could be the consequence of limited statistical power, the result may also stem from DNMTs not having large, locus-specific effects on DNA methylation under physiological conditions. Indeed, previous work suggests that due to lack of sequence specificity, targeting of DNMTs to particular sites has to be achieved by other means such as chromatin context and transcription factors (Baubec et al., 2015; Rose and Klose, 2014; Schübeler, 2015). It is plausible that DNA methylation levels are instead predominantly regulated by factors that attract or repel the DNA methylation machinery at specific locations, instead of expression levels of the machinery itself. Finally, we note that the majority of genes that were previously identified as core epigenetic factors (EpiFactor database) are histone modifiers (Medvedeva et al., 2015). This suggests that changes in DNA methylation might be secondary to altered chromatin conformation. This idea is further supported by discussed examples such as *IKZF1, BEND3* and *NLRC5*, which are thought to attract histone modifying complexes to their binding sites. These findings are in line with the notion that DNA methylation and histone modifications are linked and can be dependent on another (Cedar and Bergman, 2009).

Conceptually, our method has similarities with previous efforts that used genetic variation to infer gene expression (Gamazon et al., 2015; Gusev et al., 2016). To the best of our knowledge, these methods have not been used to investigate directed associations between gene expression and DNA methylation. In addition, our implementation has the benefit that it accounts for LD/pleiotropy among neighboring genes, which would otherwise lead to non-specific results. This is evidenced by the large number of genes that became insignificant after accounting for LD/pleiotropy among neighboring genes.

We should note limitations of this study. First, although we systematically considered LD/pleiotropy among neighboring genes, we cannot fully rule out the possibility of pleiotropy. The genetic instruments may have pleiotropic effects on unmeasured genes, but may also affect *trans* DNA methylation through other mechanisms such as interchromosomal contacts (Bonder et al., 2016). Further studies could investigate pleiotropic effects using statistical methods such as Egger regression and heterogeneity tests (Burgess et al., 2017). These methods, however, rely on multiple independent variants which are scarce for gene expression, since most predictive variants are located near the gene and are therefore often not independent because of LD. Second, although we intend to provide a genome-wide resource of genes that influence DNA methylation, we had to limit our scope to genes that had a sufficiently predictive genetic instrument. Third, the genetic instruments generally explain a relatively small proportion of the variation in expression of their corresponding gene, which results in limited power to detect effects (median R^2^ = 0.04). In addition, although correcting for nearby GIs enabled us to obtain specific associations, it further limits the power to detect effects. Finally, we note that the detection of target CpGs in this study was limited by the coverage of the Illumina 450k array, which measures about 2% of the CpGs sites in the human genome. Whole-genome bisulfite sequencing (WGBS) would provide a higher resolution, but does so at a much higher cost.

To conclude, we present a novel resource of genes for which we provide strong evidence that they influence DNA methylation levels in blood. Our results add to the increasing evidence that transcription factors are involved in shaping the methylome and we show that our resource can provide insight into the various mechanisms through which DNA methylation is regulated (e.g. post-translation modification and secondary effects of chromatin conformation). We believe our resource will guide follow-up studies into epigenetic regulation and the role of these regulatory genes in disease.

## Methods

### Cohorts

The Biobank-based Integrative Omics Study (BIOS) Consortium comprises six Dutch biobanks: Cohort on Diabetes and Atherosclerosis Maastricht (CODAM) (van Greevenbroek et al., 2011), LifeLines-DEEP (LLD) (Tigchelaar et al., 2015), Leiden Longevity Study (LLS) (Schoenmaker et al., 2006), Netherlands Twin Registry (NTR) (Boomsma et al., 2002; Willemsen et al., 2013), Rotterdam Study (RS) (Hofman et al., 2013), and Prospective ALS Study Netherlands (PAN) (Huisman et al., 2011). Data used in this study consists of 4,162 unrelated individuals for which genotype data was available. For 4,056 of these individuals DNA methylation data was available and for 3,357 individuals RNA-sequencing data was available. Genotype data, DNA methylation data, and gene expression data were measured in whole blood. In addition, sex, age, and cell counts were obtained. The Human Genotyping facility (HugeF, Erasmus MC, Rotterdam, The Netherlands, http://www.blimdna.org) generated the methylation and RNA-sequencing data.

### Genotype data

Genotype data was generated for each cohort individually. Details on the methods used can be found in the individual papers; CODAM: (van Dam et al., 2001). LLD: (Tigchelaar et al., 2015), LLS: (Deelen et al., 2014a), NTR: (Lin et al., 2016), RS: (Hofman et al., 2013), PAN: (Huisman et al., 2011). The genotype data were harmonized towards the Genome of the Netherlands (GoNL) using Genotype Harmonizer (Deelen et al., 2014b) and subsequently imputed per cohort using MaCH (Li et al., 2010) with the Haplotype Reference Consortium panel (McCarthy et al., 2016). Per cohort, SNPs with R^2^ < 0.3 and call rate < 0.95 were removed and VCFtools (Danecek et al., 2011) was used to remove SNPs with Hardy Weinberg Equilibrium p-value < 10^−4^. After merging the cohorts, SNPs with minor allele frequency < 0.01 were removed. These imputation and filtering steps resulted in 7,568,624 SNPs that passed quality control in each of the datasets.

### Gene expression data

A detailed description regarding generation and processing of the gene expression data can be found elsewhere (Zhernakova et al., 2016). Briefly, total RNA from whole blood was deprived of globin using Ambion’s GLOBIN clear kit and subsequently processed for sequencing using Illumina’s Truseq version 2 library preparation kit. Paired-end sequencing of 2×50bp was performed using Illumina’s Hiseq2000, pooling 10 samples per lane. Finally, read sets per sample were generated using CASAVA, retaining only reads passing Illumina’s Chastity Filter for further processing. Data were generated by the Human Genotyping facility (HugeF) of ErasmusMC (The Netherlands). Initial QC was performed using FastQC (v0.10.1), removal of adaptors was performed using cutadapt (v1.1) (Martin, 2011), and Sickle (v1.2) (Joshi and Fass, 2011) was used to trim low quality ends of the reads (minimum length 25, minimum quality 20). The sequencing reads were mapped to human genome (HG19) using STAR (v2.3.0e) (Dobin et al., 2013).

To avoid reference mapping bias, all GoNL SNPs (http://www.nlgenome.nl/?page_id=9) with MAF > 0.01 in the reference genome were masked with N. Read pairs with at most 8 mismatches, mapping to as most 5 positions, were used.

Gene expression quantification was determined using base counts (Zhernakova et al., 2016). The gene definitions used for quantification were based on Ensembl version 71, with the extension that regions with overlapping exons were treated as separate genes and reads mapping within these overlapping parts did not count towards expression of the normal genes.

For data analysis we used the log counts per million (CPM). We restricted the analysis to protein-coding genes and lincRNAs (long intergenic non-coding RNA) that were at least moderately expressed (median CPM => 1). This resulted in 11,475 protein-coding genes and 355 lincRNAs that were used for further analysis. To reduce the influence of possible outliers, we transformed the data using rank-based inverse normal transformation within each cohort.

### DNA methylation data

The Zymo EZ DNA methylation kit (Zymo Research, Irvine, CA, USA) was used to bisulfite-convert 500 ng of genomic DNA, and 4 μl of bisulfite-converted DNA was measured on the Illumina HumanMethylation450 array using the manufacturer’s protocol (Illumina, San Diego, CA, USA). Preprocessing and normalization of the data were done as described in the DNAmArray workflow (https://molepi.github.io/DNAmArray_workflow/). In brief, IDAT files were read using the *minfi* (Aryee et al., 2014), while sample level quality control (QC) was performed using *MethylAid* (van Iterson et al., 2014). Filtering of individual measurements was based on detection *P*-value (*P* < 0.01), number of beads available (≤ 2) or zero values for signal intensity. Normalization was done using Functional Normalization (Fortin et al., 2014) as implemented in *minfi* (Aryee et al., 2014), using five principal components extracted using the control probes for normalization. All samples or probes with more than 5% of their values missing were removed.

#### Probe filtering

Since it has been shown that the Dutch population contains population-specific variation we identified genetic variants that overlap with probes using release 5 variant data from the GoNL project (https://molgenis26.target.rug.nl/downloads/gonl_public/variants/release5/) (Francioli et al., 2014). This data contains 20.4 million SNVs and 1.1 million short INDELs (1bp-20bp) obtained from WGS data from 498 unrelated Dutch individuals. *BCFtools* was used to extract the INFO files from the GoNL VCF files (Li et al., 2009). The genomic coordinates were stored in *GRanges* format in R (Lawrence et al., 2013), for deletions we used the length of the deletion to define the start and end coordinates of the deletion. The *findOverlaps* function in the *GenomicRanges* package was used to identify variants that were located in the SBE-site for type I probes (the SBE-site coincides with the C-nucleotide in type II probes), CpG-site or within 5 bases of the 3’-end of the probe. Since not all SNPs at SBE-sites of type I probes cause a color-channel switch, only SNPs that cause a color-channel switch (A/G, G/T, C/G SNPs for reverse strand probes and C/T, C/A and C/G SNPs for forward strand probes) and INDELs overlapping the SBE were flagged for removal. A list of all SNPs and short INDELs that overlap with 450K probes is available from: https://github.com/molepi/DNAmArray.

We identified 15,724 probes that contained one or more variants with MAF > 0.01 in the SBE-site (causing a color-channel switch), CpG-site or within 5 bases of the 3’ end and excluded these probes for further analyses. In addition, we removed probes with a non-unique mapping and non-unique 3’ nested subsequences of at least 30 bases as recommended by Zhou *et al.* (Zhou et al., 2016). In total, this led to the removal of 41,674 probes. Finally, we removed all probes on the sex chromosomes.

The final dataset consisted of 4,056 samples and 428,126 probes. To reduce the influence of possible outliers, we transformed the data using rank-based inverse normal transformation within each cohort, similar to the RNAseq data.

Proper data linkage of SNP, RNA-seq and DNA methylation array data within individuals was verified using the *omicsPrint* package (van Iterson et al., 2018).

### Imputation of missing covariates

A fraction of the samples had missing data for the phenotype measures used in subsequent analyses (white blood cell proportions, age and sex).

#### Overview missing data

White blood cell counts (Neutrophils, Eosinophils, Lymphocytes, Monocytes and Basophils) were measured as part of the complete blood cell count. Complete cell count measurements were missing for 35% of the RNAseq samples and 44% of the DNA methylation samples. Reported age and sex were missing for 1.5% of the RNAseq samples and 18% of the DNA methylation samples.

#### Imputation

Since DNA methylation and RNAseq data are informative for age, sex and white blood cell composition (Abbas et al., 2009; Horvath, 2013; Houseman et al., 2012; Peters et al., 2015), we used the data to impute these variables. Missing observations were imputed separately for the RNAseq and DNA methylation data because there is incomplete overlap between the datasets. Missing observations in the measured white blood cell counts (WBCC) were imputed using the R package *pls*, as described earlier (van Iterson et al., 2017). For missing age and sex measurements, we compared the performance of the elastic-net, LASSO, ridge and pls methods. To evaluate the performance of these models, the data was randomly split into a train set (2/3) and a test set (1/3). This procedure was repeated 25 times, each time calculating the accuracy in the test set (mean squared error for age and F_1_-score for sex). The above algorithm was performed using varying numbers of input variables (50 to 10,000), where the input variables were selected based on their correlation with the outcome. The model and number of input variables that resulted in the best average accuracy in the test sets was selected to impute missing data. The average correlation between predicted and reported age in the tests sets was 0.98 for the DNA methylation data and 0.92 for the RNAseq data. Sex was almost perfectly predicted (accuracy ≈ 0.995) in both DNA methylation and RNAseq data.

### Constructing a local genetic instrument for gene expression

We constructed a genetic instrument (GI) for the expression of each gene using nearby genetic variants. We split the genotype and RNAseq data in a training set (one-third of all samples, N = 1,119) and a test set (two-thirds of all samples, N = 2,238), making sure all cohorts and both sexes were equally represented within each set. In the train set we built a GI for the expression of each gene by employing a two-step approach in which LASSO regression is used for variable selection and coefficient estimation (Tibshirani, 1996). We previously reported that LASSO performs better (BLUP, BSLMM) or similar (elastic net) compared to other methods to create GIs (Luijk et al., 2018). The number of variables chosen by LASSO is generally large and potentially includes noise variables (Meinshausen, 2007). A two-step approach can overcome this problem, where LASSO is first used for variable selection and is then used again on the selected variables for coefficient estimation. In detail, for each gene we performed the following procedure:

1. LASSO is performed in the train set to select nearby genetic variants (within the gene or within 100kb of the gene’s transcription start site (TSS) or transcription end site (TES)) that are predictive of the expression of the respective gene. 5-fold cross-validation was used to find the penalization parameter *λ* that minimizes the mean squared error (MSE).
2. LASSO is performed in the train set on the remaining genetic variants. In order to select the most parsimonious model without losing accuracy, we used the ‘one-standard error rule’ to select the largest penalization parameter *λ* that is within 1 standard error of the minimum with the constraint that at least one SNP has a nonzero coefficient (James et al., 2013). We then calculated the genetic instrument as the sum of dosages of each SNP multiplied by their effect sizes.

In both LASSO steps we included known covariates (age, sex, cohort and white blood cell composition) and the first five principal components derived from the RNAseq data in the LASSO model, because the inclusion of covariates that explain variation will generally lead to increased precision of the SNP coefficients (Burgess et al., 2011). These covariates were left unpenalized, ensuring that their coefficient is always nonzero.

We evaluated the predictive performance of the genetic instruments in the test set. We employed Analysis of Variance (ANOVA) to evaluate the added predictive power of the GI over the covariates, as reflected by the F-statistic. Genetic instruments with an F-statistic > 10 were considered valid instruments (Staiger and Stock, 1997).

### Testing for *trans* effects

Using linear regression, we tested for an association between each GI and the DNA methylation levels of CpGs *in trans* (> 10Mb). Age, sex, cohort, white blood cell composition and five principal components were included as covariates. We used the BioConductor package *bacon* to correct for inflation and/or bias in the test statistics (van Iterson et al., 2017) and corrected for multiple testing using the Bonferroni correction (8,644 × 428,126 tests, *P* < 1.4 × 10^−11^). A two-step approach was used to account for LD/pleiotropy within the obtained results (Figure S4). First, we corrected all GI-CpG pairs for nearby GIs (within 1Mb of the gene’s TSS/TES). Genes for which the corresponding GI was highly correlated with one or more neighboring GIs (*r* > 0.95) were excluded from further analyses. To prevent collinearity, we pruned the neighboring GIs that were included in the model using the *findCorrelation* function in the *caret* R package using a correlation cutoff of 0.95 (Kuhn, 2008). Second, among the GIs that remained significant, we tested for residual pleiotropic effects that were not captured by the correction for nearby GIs. For each GI we evaluated the added predictive power over the covariates and neighboring GIs on the expression corresponding to nearby significant GIs. We excluded GIs that shared target CpGs with a neighboring significant GI (at a gene-level Bonferroni level, *P <* 1.2 × 10^−7^) and were at least weakly predictive of the expression of that gene (*F* > 5).

### Enrichment analyses

Gene set enrichment were performed for GO molecular functions using DAVID (Huang et al., 2008), where all genes with a predictive GI (*F* > 10) where used as background. Fisher’s exact test was used to test for enrichment of transcription factors (Lambert et al., 2018) and epigenetic factors (Medvedeva et al., 2015). Chromatin state segments were downloaded from the Epigenomics Roadmap for all blood subtypes (Kundaje et al., 2015). CpGs were annotated to different segments based on the most frequent occurring feature in the various blood cell subtypes. Repeat sequences were downloaded from the UCSC table browser (Karolchik, 2004). Enrichment tests for chromatin state segments and repeat sequences were performed using Fisher’s exact test.

### Association with *trans* expression levels

For several examples we tested whether the target CpGs were associated with nearby gene expression and/or if the GI corresponding to the index gene was associated with the expression levels of genes near its target CpGs. We tested for an association between the target CpGs and the expression of nearby genes (<250Kb) using linear regression. Age, sex, cohort, white blood cell composition and 10 principal components (first five PCs derived from gene expression data, and first five PCs derived from DNA methylation data) were included as covariates. Similarly, to test whether the index GI was associated with the expression of genes near the target CpGs, we tested for an association between the GI and the expression of nearby genes (<250kb) using linear regression. Age, sex, cohort and white blood cell compositions were included as covariates. In both analyses we used *bacon* to correct for bias and inflation in the test-statistics and adjusted for multiple correction using the Bonferroni correction (van Iterson et al., 2017).

### URLs

Results are available at http://bbmri.researchlumc.nl/atlas/#data. Data were generated by the Human Genotyping facility (HugeF) of ErasmusMC, the Netherlands (http://www.glimDNA.org). Webpages of participating cohorts: LifeLines, http://lifelines.nl/lifelines-research/general; Leiden Longevity Study, http://www.leidenlangleven.nl/; Netherlands Twin Registry, http://www.tweelingenregister.org/; Rotterdam Studies, http://www.epib.nl/research/ergo.htm; Genetic Research in Isolated Populations program, http://www.epib.nl/research/geneticepi/research.html#gip; CODAM study, http://www.carimmaastricht.nl/; PAN study, http://www.alsonderzoek.nl/.

## Acknowledgements

This research was financially supported by BBMRI-NL, a Research Infrastructure financed by the Dutch government (NWO, numbers 184.021.007 and 184.033.111). Samples were contributed by LifeLines, the Leiden Longevity Study, the Netherlands Twin Registry (NTR), the Rotterdam Study, the Genetic Research in Isolated Populations program, the Cohort on Diabetes and Atherosclerosis Maastricht (CODAM) study and the Prospective ALS study Netherlands (PAN). We thank the participants of all afore-mentioned biobanks and acknowledge the contributions of the investigators to this study. This work was carried out on the Dutch national e-infrastructure with the support of SURF Cooperative. We acknowledge the support from the Netherlands CardioVascular Research Initiative (the Dutch Heart Foundation, Dutch Federation of University Medical Centres, the Netherlands Organisation for Health Research and Development, and the Royal Netherlands Academy of Sciences) for the GENIUS project Generating the best evidence-based pharmaceutical targets for atherosclerosis (CVON2011-19). This work was supported by The Netherlands Heart Foundation (2017T075).

## Author contributions

Conceptualization, B.T.H., E.W.v.Z., P.J.H., R.L., K.F.D., M.v.I.; Methodology, P.J.H, R.L., B.T.H., E.W.v.Z., M.v.I.; Formal Analysis, P.J.H.; Resources, W.A., A.C., D.I.B., C.M.v.D., M.M.J.v.G., J.H.V., C.W., L.F., P.A.C.t.H., R.J., J.v.M., H.M., P.E.S., BIOS Consortium; Writing—Original Draft, P.J.H.; Writing—Review and Editing, P.J.H, R.L., B.T.H., E.W.v.Z., J.H.V., L.D., M.v.I., K.F.D., R.J., J.B.J.v.M., P.A.C.t.H., M.A.I., M.M.J.v.G., D.I.B., P.E.S.; Visualization, P.J.H., B.T.H.; Supervision, B.T.H..

## Consortia

Members BIOS consortium: Bastiaan T. Heijmans, Peter A.C. ’t Hoen, Joyce van Meurs, Dorret I. Boomsma, René Pool, Jenny van Dongen, Jouke J. Hottenga, Marleen MJ van Greevenbroek, Coen D.A. Stehouwer, Carla J.H. van der Kallen, Casper G. Schalkwijk, Cisca Wijmenga, Sasha Zhernakova, Ettje F. Tigchelaar, P. Eline Slagboom, Marian Beekman, Joris Deelen, Diana van Heemst, Jan H. Veldink, Leonard H. van den Berg, Cornelia M. van Duijn, Bert A. Hofman, André G. Uitterlinden, P. Mila Jhamai, Michael Verbiest, H. Eka D. Suchiman, Marijn Verkerk, Ruud van der Breggen, Jeroen van Rooij, Nico Lakenberg, Hailiang Mei, Jan Bot, Dasha V. Zhernakova, Peter van ’t Hof, Patrick Deelen, Irene Nooren, Matthijs Moed, Martijn Vermaat, René Luijk, Marc Jan Bonder, Maarten van Iterson, Freerk van Dijk, Michiel van Galen, Wibowo Arindrarto, Szymon M. Kielbasa, Morris A. Swertz, Erik. W van Zwet, Aaron Isaacs, Rick Jansen, Lude Franke.

## REFERENCES

Abbas, A.R., Wolslegel, K., Seshasayee, D., Modrusan, Z., and Clark, H.F. (2009). Deconvolution of Blood Microarray Data Identifies Cellular Activation Patterns in Systemic Lupus Erythematosus. PLoS ONE 4, e6098.

Aryee, M.J., Jaffe, A.E., Corrada-Bravo, H., Ladd-Acosta, C., Feinberg, A.P., Hansen, K.D., and Irizarry, R.A. (2014). Minfi: a flexible and comprehensive Bioconductor package for the analysis of Infinium DNA methylation microarrays. Bioinformatics 30, 1363–1369.

Baubec, T., Colombo, D.F., Wirbelauer, C., Schmidt, J., Burger, L., Krebs, A.R., Akalin, A., and Schübeler, D. (2015). Genomic profiling of DNA methyltransferases reveals a role for DNMT3B in genic methylation. Nature 520, 243–247.

Bjornsson, H.T. (2015). The Mendelian disorders of the epigenetic machinery. Genome Res. 25, 1473–1481.

Blattler, A., and Farnham, P.J. (2013). Cross-talk between Site-specific Transcription Factors and DNA Methylation States. J. Biol. Chem. 288, 34287–34294.

Bonder, M.J., Luijk, R., Zhernakova, D.V., Moed, M., Deelen, P., Vermaat, M., van Iterson, M., van Dijk, F., van Galen, M., Bot, J., et al. (2016). Disease variants alter transcription factor levels and methylation of their binding sites. Nat. Genet.

Bonizzi, G., and Karin, M. (2004). The two NF-κB activation pathways and their role in innate and adaptive immunity. Trends Immunol. 25, 280–288.

Boomsma, D.I., Vink, J.M., Van Beijsterveldt, T.C., de Geus, E.J., Beem, A.L., Mulder, E.J., Derks, E.M., Riese, H., Willemsen, G.A., Bartels, M., et al. (2002). Netherlands Twin Register: a focus on longitudinal research. Twin Res. Hum. Genet. 5, 401–406.

Brenner, C., Deplus, R., Didelot, C., Loriot, A., Viré, E., De Smet, C., Gutierrez, A., Danovi, D., Bernard, D., Boon, T., et al. (2005). Myc represses transcription through recruitment of DNA methyltransferase corepressor. EMBO J. 24, 336–346.

Burgess, S., Thompson, S.G., and CRP CHD Genetics Collaboration (2011). Avoiding bias from weak instruments in Mendelian randomization studies. Int. J. Epidemiol. 40, 755–764.

Burgess, S., Bowden, J., Fall, T., Ingelsson, E., and Thompson, S.G. (2017). Sensitivity Analyses for Robust Causal Inference from Mendelian Randomization Analyses with Multiple Genetic Variants: Epidemiology 28, 30–42.

Cedar, H., and Bergman, Y. (2009). Linking DNA methylation and histone modification: patterns and paradigms. Nat. Rev. Genet. 10, 295–304.

van Dam, R.M., Boer, J.M., Feskens, E.J., and Seidell, J.C. (2001). Parental history of diabetes modifies the association between abdominal adiposity and hyperglycemia. Diabetes Care 24, 1454–1459.

Danecek, P., Auton, A., Abecasis, G., Albers, C.A., Banks, E., DePristo, M.A., Handsaker, R.E., Lunter, G., Marth, G.T., Sherry, S.T., et al. (2011). The variant call format and VCFtools. Bioinformatics 27, 2156–2158.

Daxinger, L., Harten, S.K., Oey, H., and Epp, T. (2013). An ENU mutagenesis screen identifies novel and known genes involved in epigenetic processes in the mouse. Genome Biol. 14.

Deelen, J., Beekman, M., Uh, H.-W., Broer, L., Ayers, K.L., Tan, Q., Kamatani, Y., Bennet, A.M., Tamm, R., Trompet, S., et al. (2014a). Genome-wide association meta-analysis of human longevity identifies a novel locus conferring survival beyond 90 years of age. Hum. Mol. Genet. 23, 4420–4432.

Deelen, P., Bonder, M.J., van der Velde, K.J., Westra, H.-J., Winder, E., Hendriksen, D., Franke, L., and Swertz, M.A. (2014b). Genotype harmonizer: automatic strand alignment and format conversion for genotype data integration. BMC Res. Notes 7, 901.

Di Croce, L., Raker, V., Corsaro, M., Fazi, F., Fanelli, M., Faretta, M., Fuks, F., Lo Coco, F., Kouzarides, T., Nervi, C., et al. (2002). Methyltransferase recruitment and DNA hypermethylation of target promoters by an oncogenic transcription factor. Science 295, 1079–1082.

Dobin, A., Davis, C.A., Schlesinger, F., Drenkow, J., Zaleski, C., Jha, S., Batut, P., Chaisson, M., and Gingeras, T.R. (2013). STAR: ultrafast universal RNA-seq aligner. Bioinformatics 29, 15–21.

Dor, Y., and Cedar, H. (2018). Principles of DNA methylation and their implications for biology and medicine. The Lancet 392, 777–786.

Filion, G.J.P., Zhenilo, S., Salozhin, S., Yamada, D., Prokhortchouk, E., and Defossez, P.-A. (2006). A Family of Human Zinc Finger Proteins That Bind Methylated DNA and Repress Transcription. Mol. Cell. Biol. 26, 169–181.

Fortin, J.-P., Labbe, A., Lemire, M., Zanke, B.W., Hudson, T.J., Fertig, E.J., Greenwood, C.M., and Hansen, K.D. (2014). Functional normalization of 450k methylation array data improves replication in large cancer studies. Genome Biol. 15, 1–17.

Francioli, L.C., Menelaou, A., Pulit, S.L., van Dijk, F., Palamara, P.F., Elbers, C.C., Neerincx, P.B.T., Ye, K., Guryev, V., Kloosterman, W.P., et al. (2014). Whole-genome sequence variation, population structure and demographic history of the Dutch population. Nat. Genet. 46, 818–825.

Gamazon, E.R., Wheeler, H.E., Shah, K.P., Mozaffari, S.V., Aquino-Michaels, K., Carroll, R.J., Eyler, A.E., Denny, J.C., Nicolae, D.L., Cox, N.J., et al. (2015). A gene-based association method for mapping traits using reference transcriptome data. Nat. Genet. 47, 1091–1098.

Garvin, A.J., Densham, R.M., Blair-Reid, S.A., Pratt, K.M., Stone, H.R., Weekes, D., Lawrence, K.J., and Morris, J.R. (2013). The deSUMOylase SENP7 promotes chromatin relaxation for homologous recombination DNA repair. EMBO Rep. 14, 975–983.

Gaunt, T.R., Shihab, H.A., Hemani, G., Min, J.L., Woodward, G., Lyttleton, O., Zheng, J., Duggirala, A., McArdle, W.L., Ho, K., et al. (2016). Systematic identification of genetic influences on methylation across the human life course. Genome Biol. 17.

van Greevenbroek, M.M.J., Jacobs, M., van der Kallen, C.J.H., Vermeulen, V.M.M.-J., Jansen, E.H.J.M., Schalkwijk, C.G., Ferreira, I., Feskens, E.J.M., and Stehouwer, C.D.A. (2011). The cross-sectional association between insulin resistance and circulating complement C3 is partly explained by plasma alanine aminotransferase, independent of central obesity and general inflammation (the CODAM study). Eur. J. Clin. Invest. 41, 372–379.

Groner, A.C., Meylan, S., Ciuffi, A., Zangger, N., Ambrosini, G., Dénervaud, N., Bucher, P., and Trono, D. (2010). KRAB–Zinc Finger Proteins and KAP1 Can Mediate Long-Range Transcriptional Repression through Heterochromatin Spreading. PLoS Genet. 6, e1000869.

Gusev, A., Ko, A., Shi, H., Bhatia, G., Chung, W., Penninx, B.W.J.H., Jansen, R., de Geus, E.J.C., Boomsma, D.I., Wright, F.A., et al. (2016). Integrative approaches for large-scale transcriptome-wide association studies. Nat. Genet. 48, 245–252.

Hayden, M.S., West, A.P., and Ghosh, S. (2006). NF-κB and the immune response. Oncogene 25, 6758–6780.

Hofman, A., Murad, S.D., van Duijn, C.M., Franco, O.H., Goedegebure, A., Arfan Ikram, M., Klaver, C.C.W., Nijsten, T.E.C., Peeters, R.P., Stricker, B.H.Ch., et al. (2013). The Rotterdam Study: 2014 objectives and design update. Eur. J. Epidemiol. 28, 889–926.

Horvath, S. (2013). DNA methylation age of human tissues and cell types. Genome Biol. 14, 1–20.

Houseman, E.A., Accomando, W.P., Koestler, D.C., Christensen, B.C., Marsit, C.J., Nelson, H.H., Wiencke, J.K., and Kelsey, K.T. (2012). DNA methylation arrays as surrogate measures of cell mixture distribution. BMC Bioinformatics 13, 1.

Huang, D.W., Sherman, B.T., and Lempicki, R.A. (2008). Systematic and integrative analysis of large gene lists using DAVID bioinformatics resources. Nat. Protoc. 4, 44–57.

Huisman, M.H.B., de Jong, S.W., van Doormaal, P.T.C., Weinreich, S.S., Schelhaas, H.J., van der Kooi, A.J., de Visser, M., Veldink, J.H., and van den Berg, L.H. (2011). Population based epidemiology of amyotrophic lateral sclerosis using capture-recapture methodology. J. Neurol. Neurosurg. Psychiatry 82, 1165–1170.

van Iterson, M., Tobi, E.W., Slieker, R.C., den Hollander, W., Luijk, R., Slagboom, P.E., and Heijmans, B.T. (2014). MethylAid: visual and interactive quality control of large Illumina 450k datasets. Bioinformatics 30, 3435–3437.

van Iterson, M., van Zwet, E.W., and Heijmans, B.T. (2017). Controlling bias and inflation in epigenome- and transcriptome-wide association studies using the empirical null distribution. Genome Biol. 18.

van Iterson, M., Cats, D., Hop, P., BIOS Consortium, and Heijmans, B.T. (2018). omicsPrint: detection of data linkage errors in multiple omics studies. Bioinformatics 34, 2142–2143.

Iyengar, S., Ivanov, A.V., Jin, V.X., Rauscher, F.J., and Farnham, P.J. (2011). Functional Analysis of KAP1 Genomic Recruitment. Mol. Cell. Biol. 31, 1833–1847.

James, G., Witten, D., Hastie, T., and Tibshirani, R. (2013). An Introduction to Statistical Learning (Springer Texts in Statistics).

Joshi, N.A., and Fass, J.N. (2011). Sickle: A sliding-window, adaptive, quality-based trimming tool for FastQ files (version 1.33).

Karolchik, D. (2004). The UCSC Table Browser data retrieval tool. Nucleic Acids Res. 32, 493D – 496.

Kobayashi, K.S., and van den Elsen, P.J. (2012). NLRC5: a key regulator of MHC class I-dependent immune responses. Nat. Rev. Immunol. 12, 813–820.

Kuhn, M. (2008). Building Predictive Models in R Using the caret Package. J. Stat. Softw. 28.

Kundaje, A., Meuleman, W., Ernst, J., Bilenky, M., Yen, A., Heravi-Moussavi, A., Kheradpour, P., Zhang, Z., Wang, J., Ziller, M.J., et al. (2015). Integrative analysis of 111 reference human epigenomes. Nature 518, 317–330.

Lambert, S.A., Jolma, A., Campitelli, L.F., Das, P.K., Yin, Y., Albu, M., Chen, X., Taipale, J., Hughes, T.R., and Weirauch, M.T. (2018). The Human Transcription Factors. Cell 172, 650–665.

Lawrence, M., Huber, W., Pagès, H., Aboyoun, P., Carlson, M., Gentleman, R., Morgan, M.T., and Carey, V.J. (2013). Software for Computing and Annotating Genomic Ranges. PLoS Comput. Biol. 9.

Lemire, M., Zaidi, S.H.E., Ban, M., Ge, B., Aïssi, D., Germain, M., Kassam, I., Wang, M., Zanke, B.W., Gagnon, F., et al. (2015). Long-range epigenetic regulation is conferred by genetic variation located at thousands of independent loci. Nat. Commun. 6, 6326.

Li, H., Handsaker, B., Wysoker, A., Fennell, T., Ruan, J., Homer, N., Marth, G., Abecasis, G., Durbin, R., and 1000 Genome Project Data Processing Subgroup (2009). The Sequence Alignment/Map format and SAMtools. Bioinformatics 25, 2078–2079.

Li, N., Johnson, D.C., Weinhold, N., Studd, J.B., Orlando, G., Mirabella, F., Mitchell, J.S., Meissner, T., Kaiser, M., Goldschmidt, H., et al. (2016). Multiple myeloma risk variant at 7p15.3 creates an IRF4-binding site and interferes with CDCA7L expression. Nat. Commun. 7, 13656.

Li, Y., Willer, C.J., Ding, J., Scheet, P., and Abecasis, G.R. (2010). MaCH: using sequence and genotype data to estimate haplotypes and unobserved genotypes. Genet. Epidemiol. 34, 816–834.

Lin, B.D., Willemsen, G., Abdellaoui, A., Bartels, M., Ehli, E.A., Davies, G.E., Boomsma, D.I., and Hottenga, J.J. (2016). The Genetic Overlap Between Hair and Eye Color. Twin Res. Hum. Genet. 19, 595–599.

Liu, T., Zhang, L., Joo, D., and Sun, S.-C. (2017). NF-κB signaling in inflammation. Signal Transduct. Target. Ther. 2, 17023.

Luijk, R., Dekkers, K.F., van Iterson, M., Arindrarto, W., Claringbould, A., Hop, P., Boomsma, D.I., van Duijn, C.M., van Greevenbroek, M.M.J., Veldink, J.H., et al. (2018). Genome-wide identification of directed gene networks using large-scale population genomics data. Nat. Commun. 9.

Lupo, A., Cesaro, E., Montano, G., Zurlo, D., Izzo, P., and Costanzo, P. (2013). KRAB-zinc finger proteins: a repressor family displaying multiple biological functions. Curr. Genomics 14, 268–278.

Marchal, C., and Miotto, B. (2015). Emerging Concept in DNA Methylation: Role of Transcription Factors in Shaping DNA Methylation Patterns: Transcription Factors in DNA Methylation. J. Cell. Physiol. 230, 743–751.

Martin, M. (2011). Cutadapt removes adapter sequences from high-throughput sequencing reads. EMBnet J. 17, pp–10.

McCarthy, S., Das, S., Kretzschmar, W., Delaneau, O., Wood, A.R., Teumer, A., Kang, H.M., Fuchsberger, C., Danecek, P., Sharp, K., et al. (2016). A reference panel of 64,976 haplotypes for genotype imputation. Nat. Genet. 48, 1279–1283.

Medvedeva, Y.A., Lennartsson, A., Ehsani, R., Kulakovskiy, I.V., Vorontsov, I.E., Panahandeh, P., Khimulya, G., Kasukawa, T., The FANTOM Consortium, and Drabløs, F. (2015). EpiFactors: a comprehensive database of human epigenetic factors and complexes. Database 2015, 1–10.

Meinshausen, N. (2007). Relaxed lasso. Comput. Stat. Data Anal. 52, 374–393.

Meylan, S., Groner, A.C., Ambrosini, G., Malani, N., Quenneville, S., Zangger, N., Kapopoulou, A., Kauzlaric, A., Rougemont, J., Ciuffi, A., et al. (2011). A gene-rich, transcriptionally active environment and the pre-deposition of repressive marks are predictive of susceptibility to KRAB/KAP1-mediated silencing. BMC Genomics 12, 378.

Mungall, A.J., Palmer, S.A., Sims, S.K., Edwards, C.A., Ashurst, J.L., Wilming, L., Jones, M.C., Horton, R., Hunt, S.E., Scott, C.E., et al. (2003). The DNA sequence and analysis of human chromosome 6. 425, 8.

Orrù, V., Steri, M., Sole, G., Sidore, C., Virdis, F., Dei, M., Lai, S., Zoledziewska, M., Busonero, F., Mulas, A., et al. (2013). Genetic Variants Regulating Immune Cell Levels in Health and Disease. Cell 155, 242–256.

Peters, M.J., Joehanes, R., Pilling, L.C., Schurmann, C., Conneely, K.N., Powell, J., Reinmaa, E., Sutphin, G.L., Zhernakova, A., Schramm, K., et al. (2015). The transcriptional landscape of age in human peripheral blood. Nat. Commun. 6, 8570.

Quenneville, S., Verde, G., Corsinotti, A., Kapopoulou, A., Jakobsson, J., Offner, S., Baglivo, I., Pedone, P.V., Grimaldi, G., Riccio, A., et al. (2011). In Embryonic Stem Cells, ZFP57/KAP1 Recognize a Methylated Hexanucleotide to Affect Chromatin and DNA Methylation of Imprinting Control Regions. Mol. Cell 44, 361–372.

de la Rica, L., Rodríguez-Ubreva, J., García, M., Islam, A.B., Urquiza, J.M., Hernando, H., Christensen, J., Helin, K., Gómez-Vaquero, C., and Ballestar, E. (2013). PU. 1 target genes undergo Tet2-coupled demethylation and DNMT3b-mediated methylation in monocyte-to-osteoclast differentiation. Genome Biol. 14, R99.

Roederer, M., Quaye, L., Mangino, M., Beddall, M.H., Mahnke, Y., Chattopadhyay, P., Tosi, I., Napolitano, L., Terranova Barberio, M., Menni, C., et al. (2015). The Genetic Architecture of the Human Immune System: A Bioresource for Autoimmunity and Disease Pathogenesis. Cell 161, 387–403.

Rose, N.R., and Klose, R.J. (2014). Understanding the relationship between DNA methylation and histone lysine methylation. Biochim. Biophys. Acta BBA - Gene Regul. Mech. 1839, 1362–1372.

Saksouk, N., Barth, T.K., Ziegler-Birling, C., Olova, N., Nowak, A., Rey, E., Mateos-Langerak, J., Urbach, S., Reik, W., Torres-Padilla, M.-E., et al. (2014). Redundant Mechanisms to Form Silent Chromatin at Pericentromeric Regions Rely on BEND3 and DNA Methylation. Mol. Cell 56, 580–594.

Schoenmaker, M., de Craen, A.J., de Meijer, P.H., Beekman, M., Blauw, G.J., Slagboom, P.E., and Westendorp, R.G. (2006). Evidence of genetic enrichment for exceptional survival using a family approach: the Leiden Longevity Study. Eur. J. Hum. Genet. EJHG 14, 79.

Schübeler, D. (2015). Function and information content of DNA methylation. Nature 517, 321–326.

Sen, P., Shah, P.P., Nativio, R., and Berger, S.L. (2016). Epigenetic Mechanisms of Longevity and Aging. Cell 166, 822–839.

Shen, H., and Laird, P.W. (2013). Interplay between the Cancer Genome and Epigenome. Cell 153, 38–55.

Stadler, M.B., Murr, R., Burger, L., Ivanek, R., Lienert, F., Schöler, A., Wirbelauer, C., Oakeley, E.J., Gaidatzis, D., Tiwari, V.K., et al. (2011). DNA-binding factors shape the mouse methylome at distal regulatory regions. Nature.

Staiger, D., and Stock, J.H. (1997). Instrumental variables regression with weak instruments. Econometrica 65, 557–586.

Thijssen, P.E., Ito, Y., Grillo, G., Wang, J., Velasco, G., Nitta, H., Unoki, M., Yoshihara, M., Suyama, M., Sun, Y., et al. (2015). Mutations in CDCA7 and HELLS cause immunodeficiency–centromeric instability–facial anomalies syndrome. Nat. Commun. 6, 7870.

Tibshirani, R. (1996). Regression Shrinkage and Selection via the Lasso. J R Stat Soc Ser B 58, 267–288.

Tigchelaar, E.F., Zhernakova, A., Dekens, J.A., Hermes, G., Baranska, A., Mujagic, Z., Swertz, M.A., Muñoz, A.M., Deelen, P., Cénit, M.C., et al. (2015). Cohort profile: LifeLines DEEP, a prospective, general population cohort study in the northern Netherlands: study design and baseline characteristics. BMJ Open 5, e006772.

Velasco, G., Hube, F., Rollin, J., Neuillet, D., Philippe, C., Bouzinba-Segard, H., Galvani, A., Viegas-Pequignot, E., and Francastel, C. (2010). Dnmt3b recruitment through E2F6 transcriptional repressor mediates germ-line gene silencing in murine somatic tissues. Proc. Natl. Acad. Sci. 107, 9281–9286.

Velasco, G., Grillo, G., Touleimat, N., Ferry, L., Ivkovic, I., Ribierre, F., Deleuze, J.-F., Chantalat, S., Picard, C., and Francastel, C. (2018). Comparative methylome analysis of ICF patients identifies heterochromatin loci that require ZBTB24, CDCA7 and HELLS for their methylated state. Hum. Mol. Genet. 27, 2409–2424.

Wagner, S., Hess, M.A., Ormonde-Hanson, P., Malandro, J., Hu, H., Chen, M., Kehrer, R., Frodsham, M., Schumacher, C., Beluch, M., et al. (2000). A Broad Role for the Zinc Finger Protein ZNF202 in Human Lipid Metabolism. J. Biol. Chem. 275, 15685–15690.

Wang, Y., Xiao, M., Chen, X., Chen, L., Xu, Y., Lv, L., Wang, P., Yang, H., Ma, S., Lin, H., et al. (2015). WT1 Recruits TET2 to Regulate Its Target Gene Expression and Suppress Leukemia Cell Proliferation. Mol. Cell 57, 662–673.

Willemsen, G., Vink, J.M., Abdellaoui, A., den Braber, A., van Beek, J.H.D.A., Draisma, H.H.M., van Dongen, J., van ‘t Ent, D., Geels, L.M., van Lien, R., et al. (2013). The Adult Netherlands Twin Register: Twenty-Five Years of Survey and Biological Data Collection. Twin Res. Hum. Genet. 16, 271–281.

Zentner, G.E., and Henikoff, S. (2013). Regulation of nucleosome dynamics by histone modifications. Nat. Struct. Mol. Biol. 20, 259–266.

Zhernakova, D.V., Deelen, P., Vermaat, M., van Iterson, M., van Galen, M., Arindrarto, W., van ‘t Hof, P., Mei, H., van Dijk, F., Westra, H.-J., et al. (2016). Identification of context-dependent expression quantitative trait loci in whole blood. Nat. Genet. 49, 139–145.

Zhou, W., Laird, P.W., and Shen, H. (2016). Comprehensive characterization, annotation and innovative use of Infinium DNA methylation BeadChip probes. Nucleic Acids Res. gkw967.

Zhu, H., Wang, G., and Qian, J. (2016). Transcription factors as readers and effectors of DNA methylation. Nat. Rev. Genet. 17, 551–565.

Zuo, X., Sheng, J., Lau, H.-T., McDonald, C.M., Andrade, M., Cullen, D.E., Bell, F.T., Iacovino, M., Kyba, M., Xu, G., et al. (2012). Zinc Finger Protein ZFP57 Requires Its Co-factor to Recruit DNA Methyltransferases and Maintains DNA Methylation Imprint in Embryonic Stem Cells via Its Transcriptional Repression Domain. J. Biol. Chem. 287, 2107–2118.

