## Supplementary Material for "Genome-wide identification of genes regulating DNA methylation using genetic anchors for causal inference"

| **Study** | **Number of overlapping CpGs** | **Number of overlapping CpGs with corresponding GI and *trans*-meQTL SNP in close proximity (<1Mb)** |
|---|---|---|
| Gaunt *et al.* (2016) | 413 | 408 |
| Lemire *et al.* (2015) | 504 | 489 |
| Total | 692 | 672 |

**Table S4**
We found a considerable overlap (N = 692) between the target CpGs identified in our study and the CpGs identified in two independent *trans*-meQTL studies (Gaunt et al., 2016; Lemire et al., 2015) (table S5-6). For the great majority of overlapping CpGs, the corresponding GI and *trans*-meQTL SNP were in close proximity (N = 672; 97%).

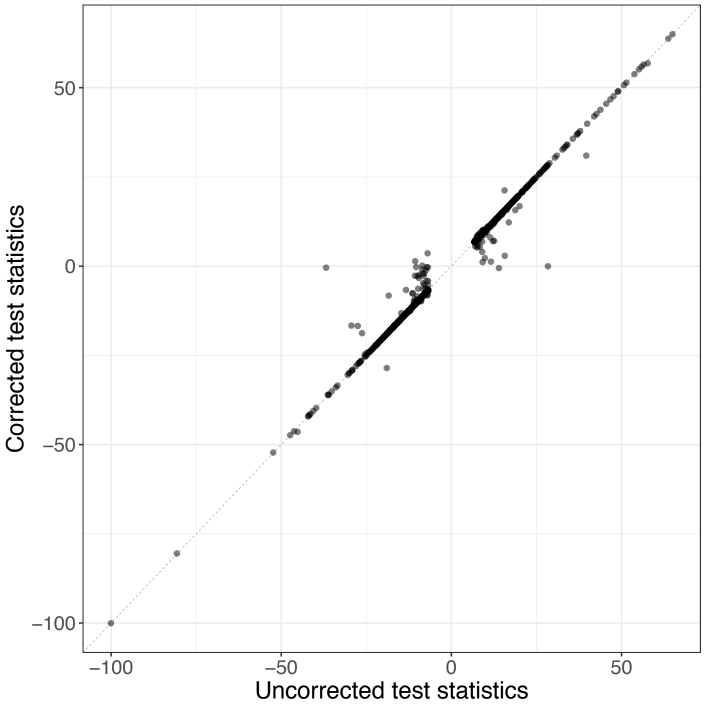


**Figure S1**

Test-statistics of the 2,633 GI-CpG pairs before (x-axis) and after (y-axis) adjustment for nearby (<1Mb) SNPs associated with white blood cell composition. 48 GI-CpG pairs were insignificant after adjustment.

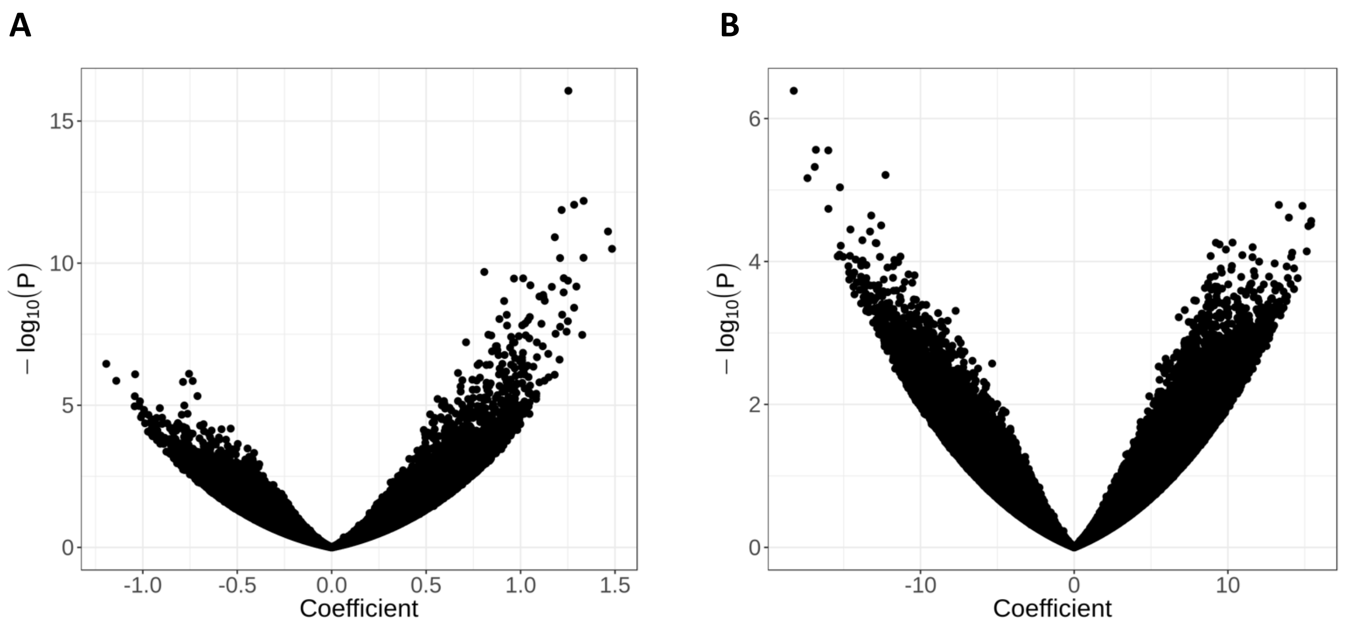


**Figure S2**

Volcano plot showing the estimated coefficients (x-axis) and –$\log_{10}$(*P*-values) (y-axis) of the genetic instrument corresponding to (a) *DNMT3A* and (b) *DNMT1* on *trans* DNA methylation levels.

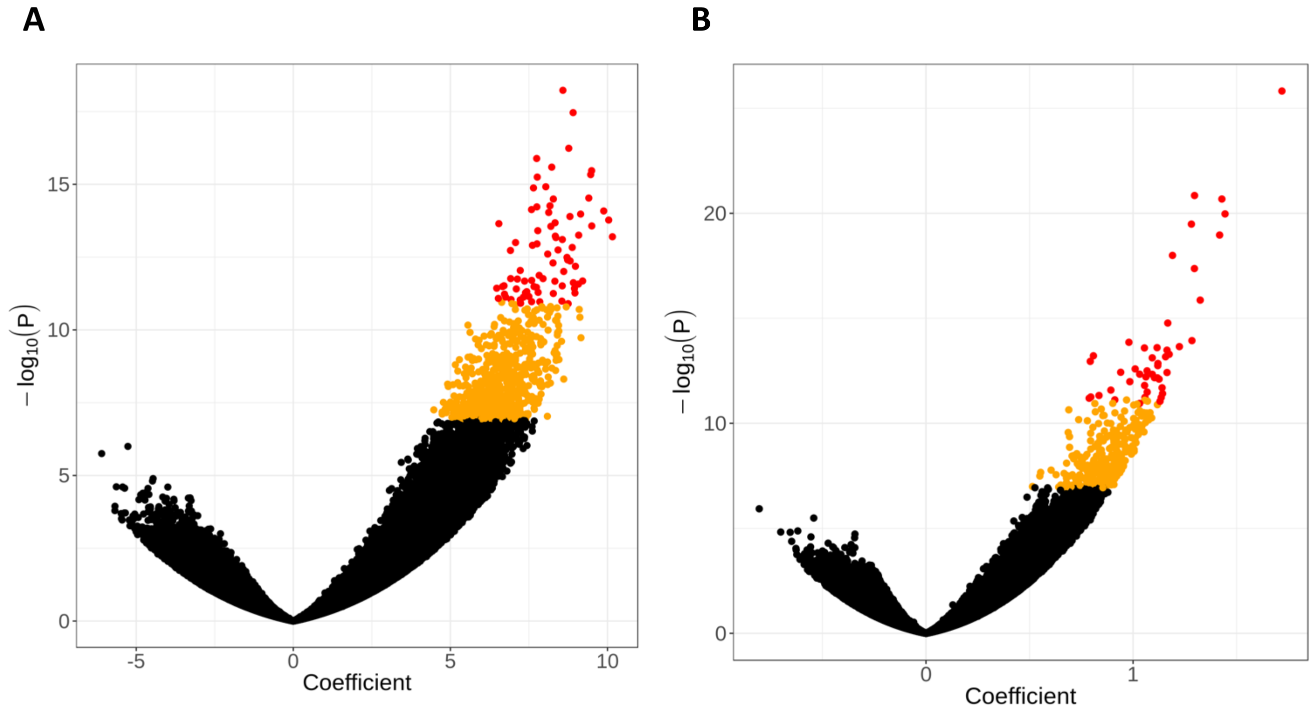


**Figure S3**

Volcano plot showing the estimated coefficients (x-axis) and $-\log_{10}$(*P*-values) (y-axis) of the genetic instrument corresponding to (a) *CDCA7* and (b) *CDCA7L* on *trans* DNA methylation levels. Red dots indicate CpG-sites that are significant at $P < 1.4 \times 10^{-11}$ and orange dots indicate CpG-sites that are significant at $P < 1.2 \times 10^{-7}$ (corresponding to a gene-level look up, 428,126 tests).

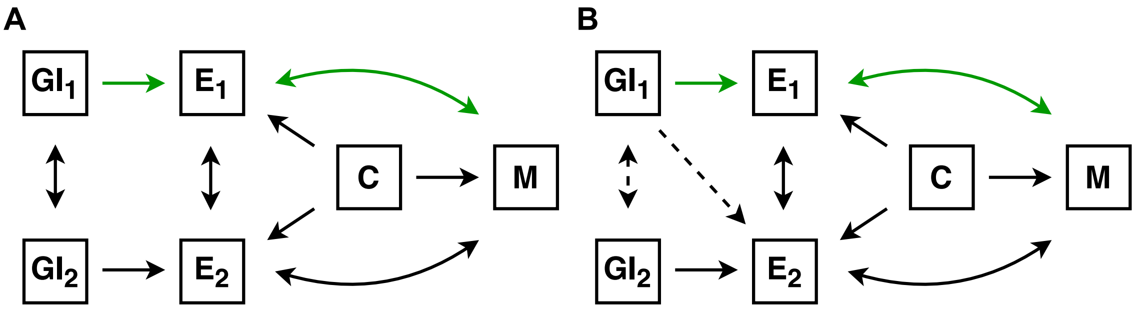


**Figure S4**

Diagrams showing the presumed relations between the genetic instruments (GI), expression (E), confounders (C) and DNA methylation (M). Single arrows indicate a causal effect; double arrows indicate that the causal effect could be in either direction. We aim to identify the effect of a GI ($GI_1$) on DNA methylation through the expression of its corresponding gene ($E_1$). Although GIs are not affected by confounding factors, they can be associated with DNA methylation through correlation with a neighboring GI ($GI_2$) and/or correlation with the expression corresponding to a neighboring GI ($E_2$) **(a)** Genetic instruments (GIs) can be associated with methylation levels through correlation with nearby GIs. To block this backdoor path, we corrected each GI for all GIs within 1Mb. **(b)** It is possible that a GI is associated with the expression of a nearby gene independent of correlation with the GI corresponding to that gene. We therefore assessed whether significant GIs that shared target CpGs were predictive of each other's gene expression, in which case they were excluded.
